# Sound localization is biased by simultaneous, and delayed by preceding visual distractors

**DOI:** 10.64898/2026.05.12.724474

**Authors:** Francesca Rocchi, Nina C. Haukes, A. John van Opstal, Marc M. van Wanrooij

## Abstract

Vision can shape auditory perception, especially when visual cues occur at different times and locations than sounds. Simultaneous but spatially misaligned lights bias the perceived location of a sound—a phenomenon known as the ventriloquism effect. Temporally misaligned lights can also affect the latency of auditory responses. However, it remains unclear how multiple visual stimuli that differ from sounds in both space and time jointly influence localization behaviour. We investigated how visual distractors, spatially misaligned by 10°, presented before and/or during a target sound influence localization accuracy and response latency in a rapid head-pointing task. Human listeners localized brief (150 ms) broadband noise bursts with an average root-mean-square error of 5° and a baseline latency of 252 ms. Simultaneous visual cues induced the ventriloquism effect, in which the perceived sound location was biased by 1.8°. Response latency also increased by 21 ms (273 ms). Preceding visual stimuli (2 s duration) did not induce a bias, but increased latency by 55 ms (307 ms). Introducing a 200 ms gap between the preceding light and the sound reduced this latency increase to 24 ms (276 ms), still not inducing a significant bias. When we presented both a preceding and a simultaneous light on opposite sides of the sound, localization reflected the bias induced by the simultaneous light (1.8°) and the latency increase induced by the preceding light (by 48 ms). These findings reveal a dissociation in audiovisual integration: preceding visual stimuli primarily influence *when* a sound is responded to (latency), while simultaneous stimuli influence *where* it is perceived (accuracy). This supports causal inference models of multisensory integration and suggests distinct underlying mechanisms for spatial and temporal processing of sounds in sensorimotor circuits.

## Introduction

Sound localization in everyday environments depends not only on acoustic cues but also on information from other senses, most notably vision. Seeing a visual stimulus at the same time as hearing a sound can influence both how quickly and how accurately we respond. Such multisensory interactions have been explained by Bayesian causal inference models. According to this idea, the brain continuously estimates whether visual and auditory cues arise from a common source, and processes them accordingly: integrate when they do, and dissociate when they do not (Fetsch & Noppeney, 2023; Körding et al., 2007; Shams & Beierholm, 2010). However, as causal inference relies on Bayesian cue-integration and requires central additional processing time, it can lead to systematic spatial biases and response delays. So far, studies have typically examined these effects with isolated pairs of one sound and one visual accessory. This simplifies interpretation, but overlooks the complexity of real-world environments, where multiple visual distractors compete for attention. The key question addressed in this paper is how multiple visual events, occurring at different times and locations relative to a sound, may jointly influence sound-localization performance. Specifically, we wondered whether preceding and simultaneously presented visual distractors interact to shape the speed and accuracy of a sound-localization response, or whether these visual cues are processed by independent mechanisms.

When a sound and a visual stimulus are aligned in space and time, localization responses are fast and accurate (Corneil et al., 2002). However, when the stimuli occur at the same location but at different times, responses slow down (Diederich & Colonius, 1987; Stevenson et al., 2012) with longer onset asynchronies producing longer delays. If the visual distractor is presented synchronously with the sound but appears at a different location, the sound-localization response shifts towards the visual distractor, which is commonly referred to as the ventriloquism effect (Alais & Burr, 2004; Hairston et al., 2003; Jack & Thurlow, 1973; Kayser et al., 2023; Körding et al., 2007). Moreover, the strength of this spatial bias decreases with increasing onset asynchrony between sound and visual distractor (Corneil et al., 2002; Kayser et al., 2023).

While the effect of visual stimulus timing has been studied extensively, it remains unclear how the timing of multiple visual distractors may affect sound localization. To our knowledge, only (Kayser et al., 2023) used both a preceding and synchronous visual distractor to investigate sound-localization accuracy. They suggested that the ventriloquism effect reflected a weighted combination of the individual effects of the distractors. However, distractors were never presented at the same location, and the effect of distractors on localization reaction times was not investigated either.

To address these gaps, the present study examines the effects of both preceding and simultaneous distractors, presented either at different or at identical locations, on the sound-localization bias and on the localization reaction times. Participants localized brief noise bursts in the horizontal plane by pointing their head as fast and as accurately as possible toward the perceived sound direction. Visual distractor V_1_ was presented long before the sound target, while distractor V_2_ occurred synchronously with the sound. Each distractor was positioned 10° to the left or to right of the perceived sound location. When both distractors were presented, they could thus be presented at the same location, or 20° apart. In addition, the distractors could be presented consecutively in time or separated by a gap. This design allowed us to study how preceding and simultaneous distractors affected sound localization.

Figure 1 conceptualizes three competing hypotheses that differ in how preceding and simultaneous distractors may interact to influence sound localisation. The *independence hypothesis (I, in Fig. 1)* proposes that the response delay caused by a preceding distractor, and the spatial bias induced by a simultaneous distractor, arise from *separate, independent* mechanisms. If true, presenting both distractors in the same trial should produce independent effects: responses should be delayed due to the preceding distractor, and localization should be biased toward the simultaneous distractor. Note that this hypothesis makes similar predictions as Bayesian causal inference (Fetsch & Noppeney, 2023; Körding et al., 2007; Shams & Beierholm, 2022). According to the *suppression hypothesis (S)*, the response delay reflects a general attempt of the sensory-motor system to suppress visual distractors. According to this model, a preceding distractor captures attention, which needs to be actively supressed. This causes a delay in the sound localisation response and reduces the influence of any subsequent visual distractor. Consequently, when both distractors are presented, responses are expected to be delayed but show little or no spatial bias - regardless of whether the distractors are spatially aligned. Moreover, the influence of the suppression is expected to weaken when there is a temporal gap between the distractors. Finally, the *adaptation hypothesis (A)* assumes that the distractor effects are spatially specific. Here, the preceding distractor affects processing only at its own location, possibly through local adaptation or spatial inhibition. If the simultaneous distractor appears at the same location, its influence is expected to be reduced, or even reversed, pushing responses away from that position, like in adaptation and inhibition-of-return effects (Klein, 2000; Posner et al., 1985; Posner & Cohen, 1984; Theeuwes & Chen, 2005; Van der Stoep et al., 2017). In this case, responses are expected to be delayed and biased only when the two distractors are spatially misaligned. Also here, effects diminish when a temporal gap separates the two distractors. By using a long-lasting preceding distractor, we biased the design toward conditions under which spatial adaptation or inhibition-of-return would be expected to be strong, ensuring that a failure to observe such effects could not be attributed to insufficient stimulation.

**Figure 1.**
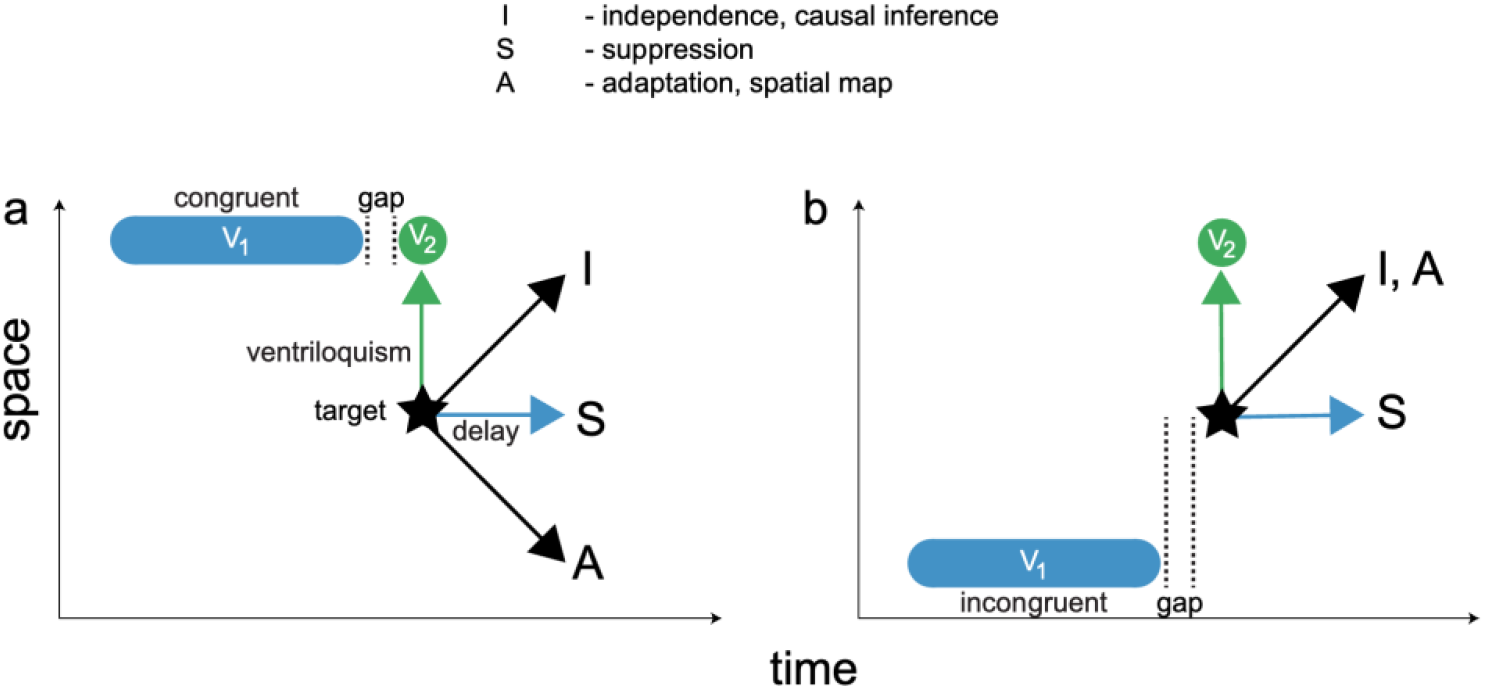
Rationale. Schematic illustration of three hypotheses (I, S, A) about how visual distractors affect the spatial accuracy (where) and response latency (when) of sound localization. The black star indicates the response to the target sound in the absence of visual distractors. The blue bar (V_1_) represents a visual distractor presented before the sound, and the green circle (V_2_) a visual distractor presented simultaneously with the sound. In the illustrated examples, V_1_ ends 200 ms before sound onset; in the experiment, this temporal gap was either 200 ms or absent (0 ms). The spatial relationship between V_1_ and V_2_ was manipulated independently: **(a)** congruent—both distractors appear at the same location, 10° away from the sound; **(b)** incongruent—distractors appear on opposite sides, each 10° from the sound. Arrows denote the predicted effects on sound-localization responses according to each hypothesis (see text, for details). The green arrow illustrates the classical ventriloquism effect: a spatial bias of perceived sound location toward a single, simultaneous visual distractor (V_2_). The blue arrow illustrates the delayed response to a sound following a single, preceding visual distractor (V_1_). The blue and black arrows indicate the predicted combined effects under the independence (I), suppression (S) and the adaptation hypotheses (A).

Our experimental design dissociates these hypotheses by varying whether a preceding and a simultaneous visual distractor are spatially aligned or opposed, while holding their spatial disparity to the sound constant. By observing whether delays and biases occur together, depend on spatial alignment, or interfere with one another, we can infer whether the underlying mechanisms operate independently or interactively. We find that preceding visual distractors robustly delay sound-localization responses without inducing spatial bias, whereas simultaneous distractors bias perceived sound location with only minor effects on response latency. When both distractors are present, their effects combine additively: localization accuracy reflects the influence of the simultaneous distractor, while response latency reflects the influence of the preceding distractor, with little evidence for mutual interference or interaction.

## Methods

### Ethics

The experiments were carried out in accordance with the relevant institutional and national regulations and with the World Medical Association Helsinki Declaration as revised in October 2013. All experimental procedures have been approved by the local ethics committee of the Faculty of Social Sciences of the Radboud University (ECSW 2016-2208-41) and participants gave their written informed consent.

### Human participants information

Eleven adult volunteers (mean age 40, 8 male, 3 female) participated in this study. They all had normal hearing as assessed by a pure-tone audiogram (within 20 dB of audiometric zero) and had normal or corrected-to-normal sight.

### Experimental setup

The experiment was conducted in a sound-attenuated room (Van Bentum et al., 2017). Participants sat at the centre of a spherical frame (radius 1.5 m) equipped with small broad-range loudspeakers ranging from −45 to 45° in the horizontal plane (negative values indicate leftward and positive values indicate rightward positions relative to the participant), at 5° increments and with light emitting diodes attached to the centre of each speaker. Head movements were recorded using the magnetic search-coil technique. Participants wore a spectacles frame (without lenses) with a mounted magnetic search-coil and a forward-facing red laser point. The laser point was used to fixate gaze, ensuring head movements were the primary means of localizing stimuli.

### Paradigm

#### auditory stimuli

The target sound was a 150 ms Gaussian white noise, band-passed between 500 and 20,000 Hz, presented at an A-weighted sound pressure level of 55 dB, including 5-ms raised-cosine onset and offset ramps. The sound was pre-generated offline in MATLAB. Visual distractors were green LEDs (wavelength 565 nm, luminance 1.4 cd/m^2^), presented before or simultaneous with the sound. Preceding distractors (V_1_) lasted 2000 ms, while simultaneous distractors (V_2_) matched the 150 ms duration of the sound target.

### Baseline Sound Localization

Prior to the main experiment, participants performed a baseline sound-localization task in which they localized brief sounds in the absence of visual distractors. This task was used to estimate each participant’s perceived sound location as a function of target position (Fig. 2a; Eq. 1), which served both to determine the individualized placement of visual distractors and to provide a reference measure of sound-only localization performance. Because the task was identical in structure to sound-only trials in the main experiment, baseline responses were later pooled with sound-only data for analysis.

**Figure 2.**
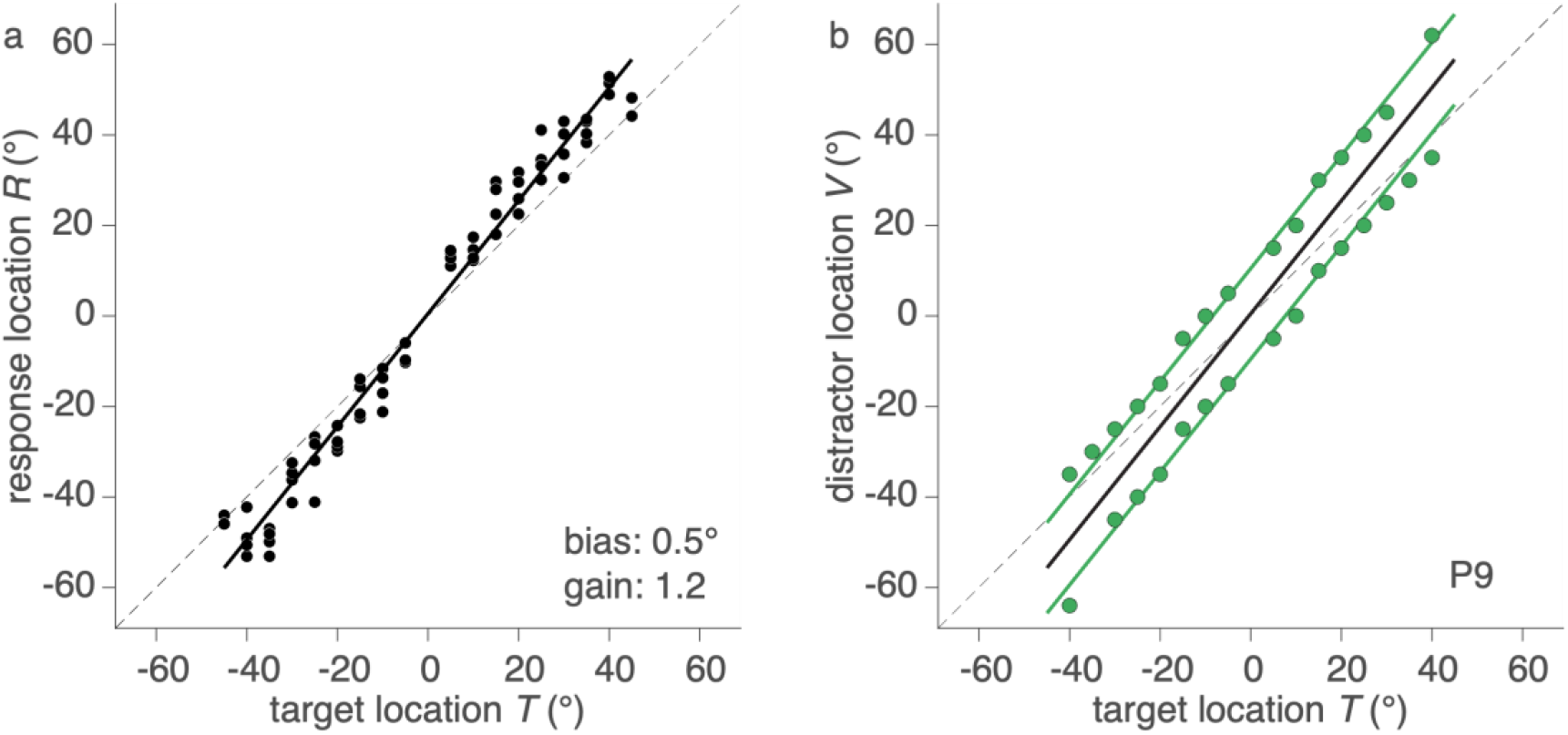
Sound localization and visual stimulus positioning. Visual distractor locations for the main experiment were customized for each participant based on performance in the baseline sound-localization task. **(a)** Stimulus–response relationship used to estimate the perceived sound location for participant P9 in the baseline sound-localization task. Black filled circles represent individual responses; the black solid line shows the linear fit to the data (gain = 1.2, bias = 0.5°). **(b)** Visual distractor locations derived from the baseline fit (black solid line, same as in (a)). LED positions with a ±10° spatial disparity relative to this fit were defined as ideal distractor locations; green filled circles indicate the actual LED positions used in the experiment (i.e., the closest available positions in the laboratory).

#### Visual stimuli

The visual distractors were positioned 10° left or right of each participant’s perceived sound location. The perceived sound location (*R*) was determined individually from a baseline sound-localization task conducted prior to the main experiment, and estimated using linear regression (Eq. 1; Fig. 2, black line):

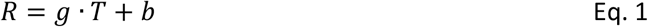

where *g*-is a dimensionless slope (gain), *b*-is an offset (bias) in degrees, and *T*-represents the location of the sound target in degrees with respect to straight-ahead. Thus, the visual distractors were positioned according to:

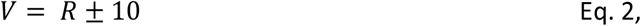

where *V*-represents the visual distractor’s horizontal position (in °). The LEDs (Fig. 2b, green circles) closest to Eq. 2 (Fig. 2b, green) were selected for the main experiment.

### Main Experiment

During the main experiment, participants localized sounds under a set of predefined stimulus configurations (Table 1). Figure 3 provides a schematic of the temporal structure of a trial for representative configurations, illustrating the order and duration of fixation, preceding distractor, gap, and target epochs. On each trial, participants fixated a central LED and pressed a button to initiate the trial. The fixation light then disappeared for 500 ms, after which either a preceding visual distractor (V_1_), the auditory target, a simultaneous visual distractor (V_2_), or a combination of these stimuli was presented, depending on the block. Participants were instructed to localize the target sound as quickly and as accurately as possible with a goal-directed head movement.

**Table 1.** Stimulus configurations per block. Overview of the stimulus configurations used in each experimental block. An “x” indicates that a given stimulus component was present. In block VI, in the case of spatially congruent distractors presented without a temporal gap, the visual stimulus formed a single continuous distractor (V_12_) rather than two separate events. V_12_ denotes a visual stimulus extending from 2000 ms before sound onset to sound offset (see text for details).

| configuration<br>block | sound | $V_1$ | $V_2$ | $V_{12}$ | gap | spatial alignment |
| --- | --- | --- | --- | --- | --- | --- |
| baseline | x | – | – | – | – | – |
| I | x | – | – | – | – | – |
| II | x | x | – | – | 200 ms | – |
| III | x | x | – | – | 0 ms | – |
| IV | x | – | x | – | – | – |
| V | x | x | x | – | 200 ms | congruent / incongruent |
| VI | x | x | x | – | 0 ms | incongruent |
| VI | x | – | – | x | 0 ms | congruent |

**Figure 3.**
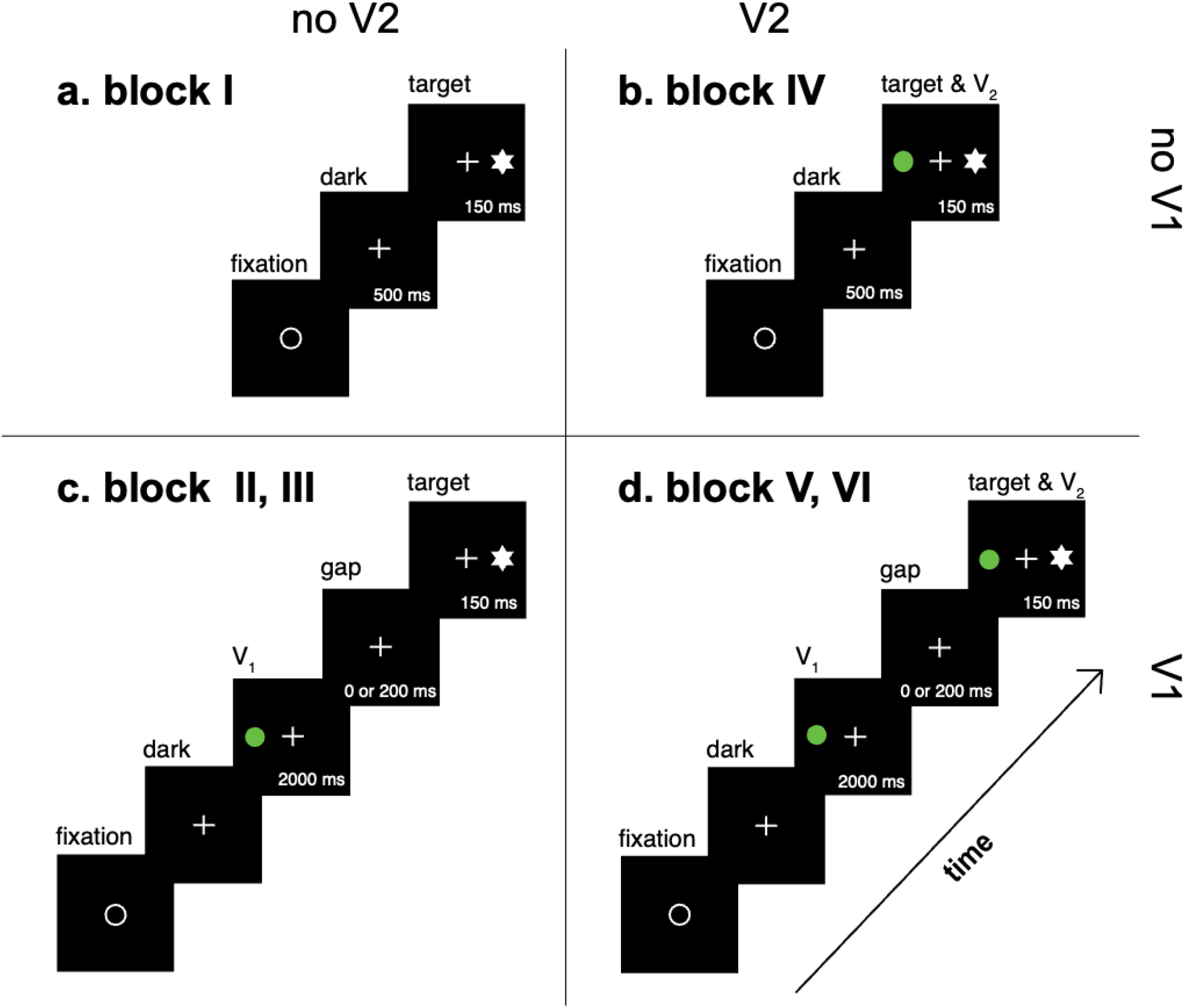
Temporal structure of representative stimulus configurations. Schematic timeline illustrating the sequence and duration of trial epochs for four representative conditions from the main experiment (see Table 1 for a complete overview of stimulus configurations). Each panel shows the order of fixation, darkness, visual stimulation, gap, and target presentation. Participants first fixated a central LED (white open circle) and initiated each trial by pressing a button, which extinguished the fixation light and was followed by 500 ms of darkness. The auditory target (white star) was then presented either alone or in combination with visual distractors, depending on the condition. Time progresses from bottom left to top right. **(a)** Sound-only condition (block I). **(b)** Simultaneous condition, in which a visual distractor (V_2_; green circle) was presented synchronously with the sound (block IV). **(c)** Preceding condition, in which a visual distractor (V_1_) preceded the sound and was offset either immediately before sound onset (“no gap”; block III) or 200 ms earlier (block II). **(d)** Two-distractor condition, in which a preceding (V_1_) and a simultaneous (V_2_) distractor were presented. Distractors could be spatially congruent (same side, shown here), or incongruent (opposite sides; ±10° from the sound). With a temporal gap this corresponds to block V; without a gap to block VI.

In the baseline block and block I, only sounds were presented. In block II, a preceding visual distractor (V_1_), starting 2200 ms before target sound onset, was presented for 2000 ms followed by a 200 ms gap. In block III, V_1_ was presented without a gap, starting 2000 ms before target sound onset. In block IV, a simultaneous visual distractor (V_2_; 150 ms) was presented synchronously with the target sound (i.e., with identical onset and offset). In block V, both a preceding and a simultaneous distractor (V_1_ and V_2_) were presented, separated by a 200 ms gap; these distractors could be either spatially congruent or incongruent (20° apart; Table 1, Fig. 3d). In block VI, the two distractors were presented without a gap. When spatially incongruent, this condition consisted of two distinct distractors (V_1_ and V_2_) at opposite locations. When spatially congruent, however, the two distractors merged into a single continuous visual stimulus extending from 2000 ms before sound onset to sound offset. This configuration is therefore treated as a single preceding distractor and is referred to as V_12_ (Table 1). Blocks I-VI were presented in random order. Within each block, target sound locations and the spatial alignment of the visual distractors were randomized across trials.

### Data analysis

All data were analysed in MATLAB (version 9.11, R2021b; Mathworks Inc., Natick, MA, USA).

#### Pre-processing

A calibration session was performed to calibrate the search-coil voltage signals into horizontal angles (described in Van Bentum et al., 2017), leading to an accuracy of better than 1°. Head saccade on- and offsets were detected using a velocity criterion of 30 °/s (see Bremen et al., 2010, for details). This procedure yielded, for each trial, a localization response defined by the movement endpoint, and a reaction time defined as the interval between sound onset and head-movement onset. Responses with reaction times shorter than 100 ms were excluded from further analysis, as they were considered predictive rather than stimulus driven. Data from the baseline block and block I were combined as both involved sound-only localisation without visual distractors.

For each participant, localization responses (Y) in all experimental blocks were corrected for individual sound-localization gain (g) and bias (b), estimated from the baseline sound-localization data (R; Eq. 1). This normalization removed idiosyncratic differences in localisation scaling and offset across participants, thereby expressing all responses in a common, participant-independent spatial reference frame. Corrected responses were obtained as:

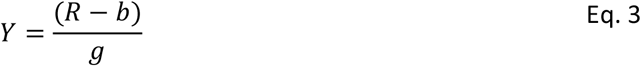

The primary outcome measures of interest were spatial accuracy and response latency. Spatial accuracy was quantified as localization error, defined as the angular difference (in °) between the target sound location and the participant’s corrected localization response. Response latency was quantified as the reciprocal of reaction time (promptness; s^−1^). This was done for statistical analysis, as promptness typically follows an approximately Gaussian distribution. Together, these measures capture the spatial (“where”) and temporal (“when”) components of sound-localization behavior.

The signs of negative V_2_ disparities (−10°) and their corresponding localization errors were inverted (i.e., multiplied by −1). In Blocks II and III (Table 1), where V_2_ was absent, the signs of negative V_1_ disparities (−10°) and their corresponding errors were inverted instead. This transformation was necessary because distractors were presented at −10° and +10°, but we are not interested in the direction of the distractor. For single-distractor conditions (V_1_, V_2_, V_12_), a positive localization error indicates that perceived sound localization was biased toward the current distractor. For the two-distractor conditions (V_1_V_2_), a positive localization error indicates that perceived sound location was biased toward the simultaneous distractor V_2_, and in incongruent trials a negative effect would mean a bias toward preceding distractor V_1_. To summarize, positive errors reflect bias toward V_2_, except in the V_1_ condition, where they reflect bias toward V_1_.

### Statistical Model

To quantify how spatial bias (localization error) and response timing (promptness) depended on the experimental manipulations, we fitted hierarchical Bayesian generalized linear models. For each dependent variable (error and promptness), the expected response was modeled as a linear combination of the experimental factors:

- Condition, C (8 levels; as discussed in Table 1),
- Target location, T (19 levels),
- Participant, P (11 levels)

including an interaction between participant and condition.

Formally, the expected response μ was modeled as:

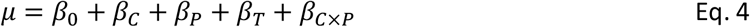

Where β_o_ represents the grand mean across all observations, and β_C_, β_P_, β_T_ capture the effects of the respective condition, participant-specific deviations, and systematic differences across target locations. All effects β were parameterized using sum-to-zero constraints, such that coefficients represent deviations from the grand mean. Participant and target-location terms were modeled as random effects (hierarchical deviations from the grand mean), absorbing variance attributable to individual differences and spatial anisotropies. These terms were included to improve model fit and generalizability but were not directly interpreted, as they are not central to testing the theoretical hypotheses.

Localization error and promptness were modeled separately but with identical factor structures. Because reaction-time derived measures and localization responses may contain occasional outliers, we modeled responses using a Student-t likelihood:

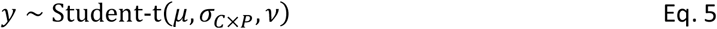

where σ_*c*×*P*-_denotes a response standard deviation determined per participant and condition, and *v* the degrees of freedom shared across conditions. The t-distribution increases robustness by allowing heavier tails without inflating condition-level variance estimates. Model adequacy was assessed using posterior predictive checks.

### Bayesian Statistical Analysis

Parameters were estimated using Bayesian inference, which provides full posterior distributions for all effects and naturally accommodates hierarchical structure. All regression coefficients were assigned flat normal priors centered at zero reflecting the expectation that most effects are small but allowing substantial deviations. The standard deviation parameter was assigned weakly informative half-normal priors. The degrees-of-freedom parameter *v* was estimated from the data with a broad prior, allowing the likelihood to range from approximately Gaussian to heavy-tailed. A hierarchical structure was used to model participant-level variability. Specifically, condition effects were partially pooled across participants, improving stability of estimates while retaining individual differences. This approach reduces overfitting and is particularly advantageous in factorial designs with some relatively sparse cells.

Posterior distributions were obtained via Markov Chain Monte Carlo sampling implemented in MATLAB (R2021b, MathWorks) using the matjags interface to JAGS (Plummer, 2003). Convergence was verified using the Gelman–Rubin statistic (all 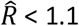) (Gelman & Rubin, 1992) and effective sample sizes exceeding 1000 for all reported parameters. Model fit was evaluated using posterior predictive checks comparing simulated data to observed distributions.

### Estimation statistics

Results are reported as posterior means together with 95% highest density intervals (HDIs). The posterior mean represents the expected value of a parameter given the data and model. The 95% HDI is the narrowest interval containing 95% of the posterior probability mass, such that any value inside the interval has higher probability density than any value outside it. An effect was considered credibly different from zero when its 95% HDI did not include zero. Because inference is based on the full posterior distribution rather than null-hypothesis testing, no p-values are reported.

## Results

### Spatial accuracy

In the absence of visual distractors, sound localization was accurate. Following bias correction (Eq. 3), the mean signed error was necessarily centered at zero (Fig. 4a; sound: 0° [−0.4, 0.4]); sampling led to a small residual deviation of 0.19° which reflects MCMC sampling variability rather than true bias, which all errors were corrected for). Introducing a simultaneous visual distractor shifted perceived sound location toward the distractor, producing a robust ventriloquism effect of approximately 1.9° (Fig. 4a, V_2_: [1.3, 2.4]°). This spatial bias was largely independent of the presence of an additional, spatially incongruent preceding distractor (Fig. 4a, V_1_V_2_ IC). When two distractors were spatially congruent and separated by a gap, the bias was slightly reduced but remained clearly positive (Fig. 4a, V_1_V_2_ C: 1.4° [0.9, 1.9]). In contrast, a preceding visual distractor alone did not bias sound localization (Fig. 4a, V_1_ and V_12_: maximum of 0.6° [0.0, 1.1]; the paired mean difference between pure sound localization and this condition includes 0), regardless of whether a temporal gap was present (V_1_) or whether the visual stimulus extended into the sound target (V_12_). Together, these results show that spatial biases in sound localization are induced by simultaneous, but not preceding, visual distractors.

**Figure 4.**
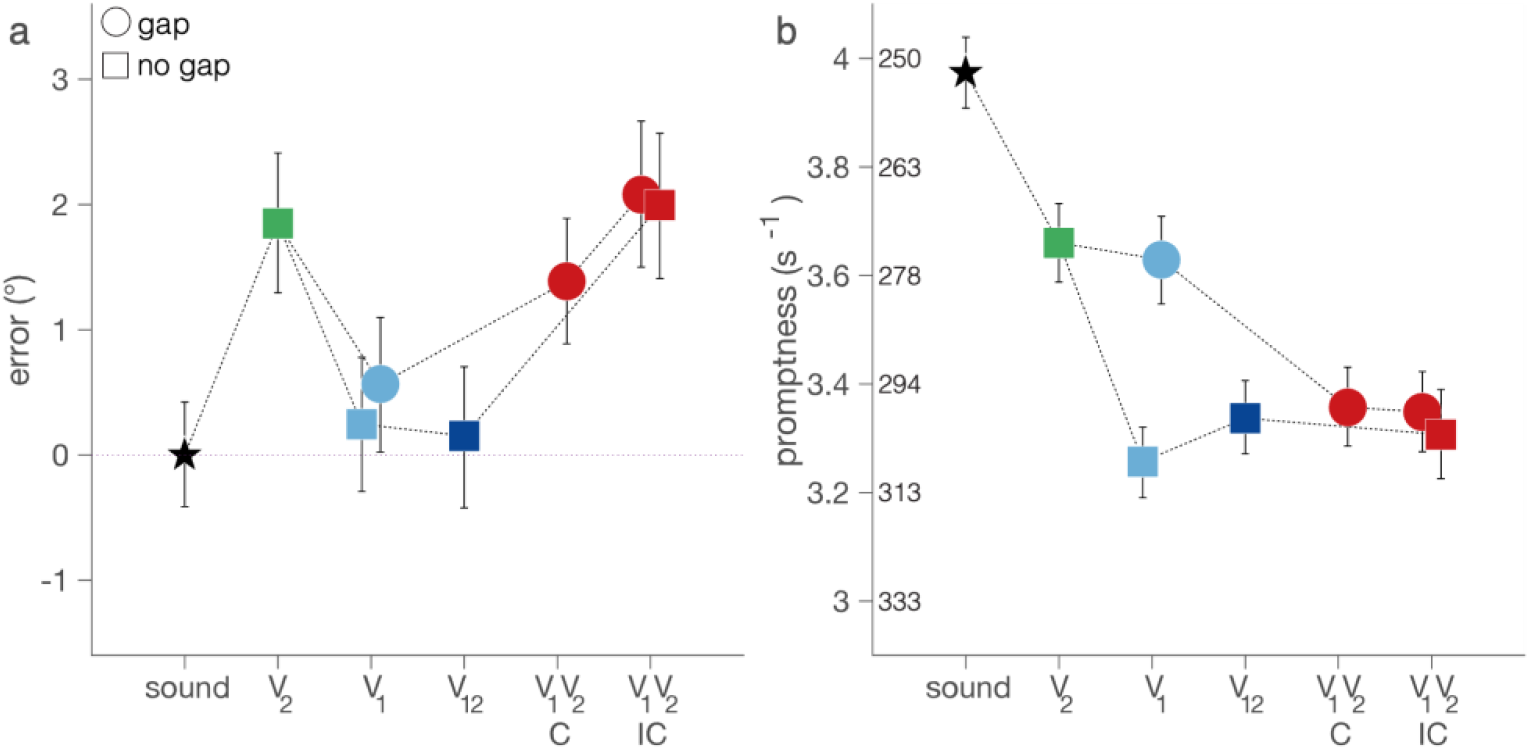
Effects of preceding (V1) and simultaneous (V2) visual distractors on spatial accuracy and` response latency. **(a)** Mean localization error across stimulus configurations. Symbols indicate sound-only trials (black star), single simultaneous distractor (green; V_2_), single preceding distractor (blue; V_1_), extended preceding distractor (V_12_), and combined distractor conditions (red; V_1_V_2_). Square symbols denote trials without a temporal gap between V_1_ and sound onset; circles denote trials with a 200 ms gap. Error bars indicate 95% highest density intervals (HDIs). **(b)** Mean response promptness across the same stimulus configurations. Values to the right of the ordinate indicate corresponding reaction times (ms).

### Response latency

In the absence of visual distractors, sound localization responses were fast, with mean reaction times around 252 ms (Fig. 4b, sound: promptness 3.97 s^−1^ [3.91, 4.04]). A single simultaneous visual distractor produced only a modest delay in responding (Fig. 4b, V_2_: 3.66 s^−1^ [3.59, 3.73]). In contrast, a preceding visual distractor substantially delayed response. When the sound followed immediately after the distractor, promptness was markedly reduced (Fig. 4b, V_1_, no gap: 3.26 s^−1^ [3.19, 3.32]). Introducing a 200 ms gap diminished, but did not eliminate, this delay (Fig. 4b, V_1_, gap: 3.63 s^−1^ [3.55, 3.71]). When both preceding and simultaneous distractors were present, response latencies were comparable to, though slightly shorter than, those observed for the preceding distractor alone (Fig. 4b, V_1_V_2_C&IC: 3.34 s^−1^ [3.25, 3.42]). Thus, response latency was primarily determined by the presence and timing of a preceding visual distractor, with little additional influence from a simultaneous distractor.

### Localization error versus delay

Figure 5 summarizes the joint effects of visual distractors on spatial accuracy and response latency by plotting localization error against response delay, both expressed relative to sound-only performance (0°, 0 ms). In this two-dimensional representation, the contrasting influences of preceding and simultaneous visual distractors become directly comparable. Across both panels, a single simultaneous distractor (V_2_) displaced responses primarily along the spatial (vertical) axis, reflecting a ventriloquism bias toward the visual stimulus, with only a small accompanying delay (green symbols). In contrast, a single preceding distractor (V_1_ or V_12_) shifted responses predominantly along the temporal axis, indicating a substantial response delay with little or no spatial bias (blue symbols). These single-distractor conditions are identical in Fig. 5a and Fig. 5b, as they do not depend on the relative spatial alignment of two distractors.

**Figure 5.**
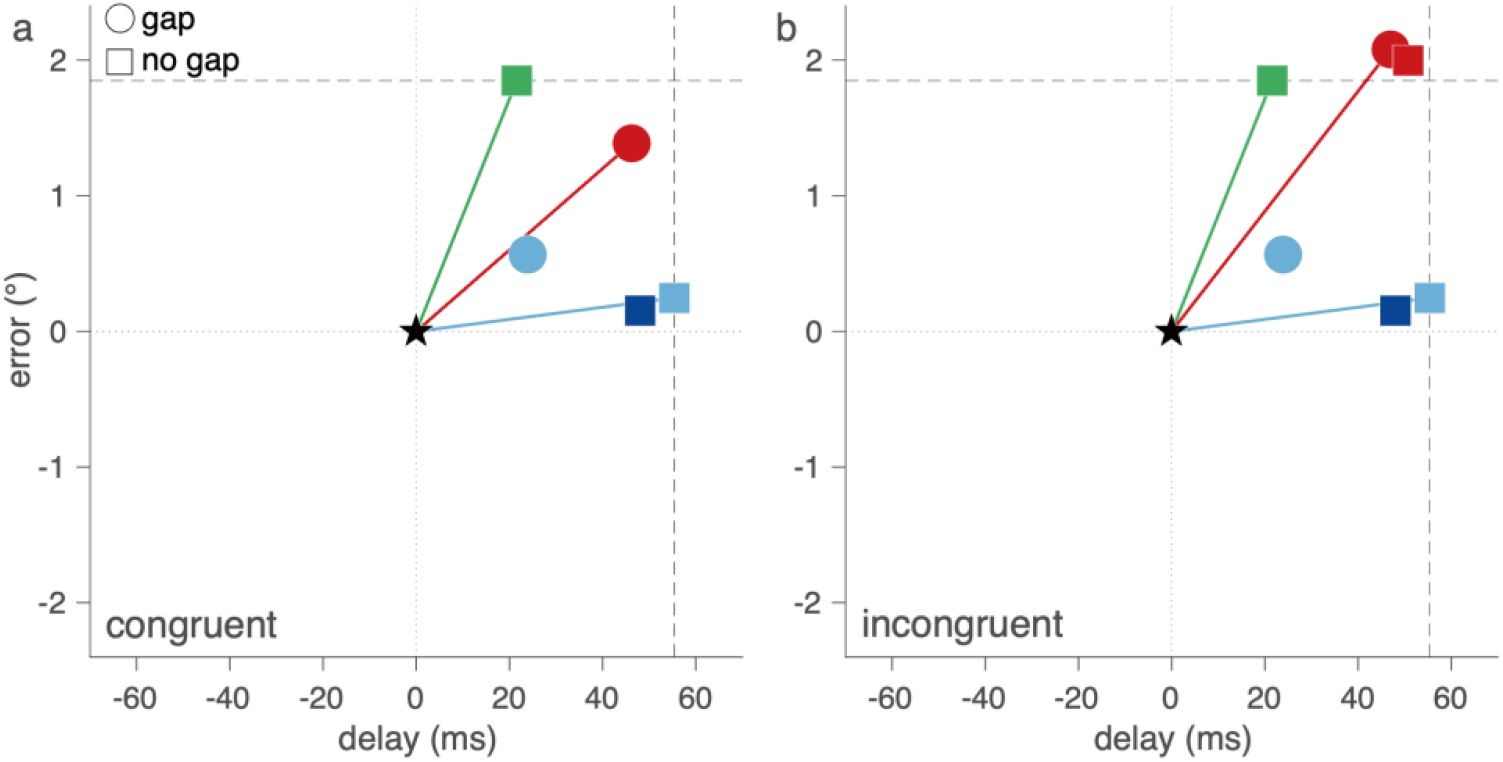
Joint effects of visual distractors on spatial accuracy and response latency. Localization error (vertical axis, in °) is plotted against response delay relative to sound-only trials (horizontal axis, in ms). The sound-only condition is shown as a black star at the origin (0°, 0 ms). Data are replotted from Fig. 4 to visualize spatial and temporal effects in a single coordinate frame. Panel **(a)** shows conditions in which preceding and simultaneous distractors were spatially congruent; panel **(b)** shows spatially incongruent conditions. Single-distractor conditions (preceding distractors V_1_/V_12_ in blue; simultaneous distractors V_2_ in green) are identical in both panels. Combined distractor conditions (V_1_V_2_) are shown in red and differ between panels depending on spatial alignment. Different marker shapes indicate the presence (circle) or absence (square) of a temporal gap between the preceding distractor and the sound.

When the preceding and simultaneous distractors were spatially congruent (Fig. 5a), combined responses occupied a region characterized by a clear spatial bias together with an increased response delay (red symbol). When the distractors were spatially incongruent (Fig. 5b), responses showed a comparable pattern: localization was biased in the direction of the simultaneous distractor, while response latency matched that induced by the preceding distractor. Thus, preceding and simultaneous distractors exert qualitatively different and largely orthogonal influences on sound-localization behavior: one primarily affecting when responses are initiated, the other affecting where sounds are perceived. The close correspondence between the predicted patterns of hypothesis I in Fig. 1 and the empirical summary in Fig. 5 underscores the additive nature of these effects, consistent with largely independent processing of spatial and temporal influences.

## Discussion

### General Findings

This study examined how visual distractors presented either before or simultaneously with a sound influence sound-localization accuracy and response latency. By independently manipulating the spatial arrangement and temporal structure of two distractors, we tested three competing accounts (Fig. 1): (I) independence of temporal and spatial influences, (S) suppression due to attentional capture by the preceding cue, and (A) spatially specific adaptation or inhibition-of-return. Across all manipulations, the data reveal a striking dissociation:

- Simultaneous distractors shift the perceived sound location (ventriloquism effect, Fig. 4a, Fig. 5), but only weakly delay responses (Fig. 4b, Fig. 5).
- Preceding distractors strongly delay responses (Fig. 4b, Fig. 5), but do not bias sound localization (Fig. 4a, Fig. 5).
- When both are presented, their effects add with minimal interaction: responses are delayed as in the preceding-cue condition, and localization is biased as in the simultaneous-cue condition (Fig. 4, Fig. 5).

This double dissociation provides strong behavioral evidence that the “where” and “when” components of multisensory orienting rely on functionally separate mechanisms.

### Spatial effects: ventriloquism depends on temporal coincidence

Our spatial results replicate classical findings in multisensory integration: when a visual stimulus occurs at a different location but at approximately the same time as the sound, localization shifts toward the visual cue (Fig. 4a; Alais & Burr, 2004; Jack & Thurlow, 1973; Körding et al., 2007; Slutsky & Recanzone, 2001; Wozny & Shams, 2011). The magnitude of this ventriloquism bias (~2°) is small compared to prior studies using similar spatial disparities and brief audiovisual targets.

The absence of a ventriloquism bias for preceding distractors (Fig. 4a) confirms the temporal constraints observed in earlier work (Kayser et al., 2023; Lewald & Guski, 2003; Van Wanrooij et al., 2009; Wallace et al., 2004): once audiovisual onsets are temporally decorrelated, the visual cue no longer contributes to the sound’s perceived location. In our design, the preceding cue was both long (2 s) and clearly separated in time from the target. This temporal mismatch appears sufficient to prevent any spatial integration.

When two distractors were presented on opposite sides, localization was always biased toward the simultaneous distractor, irrespective of whether a preceding cue was present. When both distractors occurred at the same location, the simultaneous cue still exerted its typical ventriloquism effect. Together, these findings show that spatial integration is tightly tied to temporal coincidence, supporting classical models of audiovisual binding windows (Colonius & Diederich, 2004; Slutsky & Recanzone, 2001).

### Temporal effects: preceding distraction delays orienting

Preceding distractors produced large and systematic increases in reaction time (Fig. 4b), even though they did not alter localization accuracy. The delay was strongest when the sound followed immediately after the preceding light and was substantially reduced when a 200 ms gap separated the two. This pattern mirrors the classic gap effect, in which removing a fixation stimulus facilitates the disengagement of fixation-related neural activity (Dorris & Munoz, 1995; Saslow, 1967). As such, this resembles the well-established fixation and attentional capture phenomena reported in the oculomotor literature (Corneil & Munoz, 2014; Dorris & Munoz, 1995; Goldring et al., 1996; Munoz & Corneil, 1995; Saslow, 1967). A salient visual stimulus, when presented shortly before a required orienting movement, activates the gaze fixation system and increases the time needed to initiate a movement toward a subsequent target. Thus, the temporal effects observed may not arise from crossmodal sensory integration. Instead, they may reflect fixation-related inhibition of movement initiation and disengagement dynamics in the oculomotor system carried over to head-orienting behavior. These temporal effects do not depend on the spatial location of the preceding cue and do not interact with the simultaneous distractor, further strengthening the claim that temporal and spatial influences arise from separate mechanisms.

Contrasting sharply with the large delays induced by preceding distractors, single simultaneous visual distractors produced only a modest delay in sound-localization responses. This limited increase in reaction times is consistent with earlier work showing that when multiple stimuli are presented simultaneously with small spatial disparities, reaction times increase only slightly, reflecting competition rather than disruption of movement initiation. For example, studies using double-spot visual stimuli have shown that closely spaced simultaneous targets introduce minor delays due to response selection (Ottes et al., 1984, 1985). In such cases, the oculomotor system resolves spatial competition within an already released movement state, leading to small latency costs. Our findings suggest a similar mechanism: a simultaneous visual distractor slightly delays sound localization, perhaps due to spatial competition at a sensorimotor selection stage.

### Two distractors: evidence for independence

Critically, when preceding and simultaneous distractors were combined, their effects were additive and independent (Fig. 5). A spatial bias nearly equalled the bias from a simultaneous distractor alone, and a reaction-time delay equalled the delay from a preceding distractor alone This additive pattern contradicts two hypotheses (Fig. 1, S and A). The findings are evidence against the suppression hypothesis (A). If the preceding cue triggered a global suppression of visual input (e.g., a “distractor-suppression” state), we should have seen reduced ventriloquism from the simultaneous cue, particularly in the no-gap condition. This did not occur. Spatial biases remained intact whenever the simultaneous distractor was present and temporally aligned with the sound.

The findings are also evidence against the adaptation hypothesis (A). If the preceding distractor induced location-specific adaptation, then the spatial bias should be weakened or reversed when both distractors share the same location, especially with short stimulus onset asynchronies. Again, this did not occur, except in the special case of the continuous (no-gap) co-located distractor (V_12_), which is better explained by the temporal structure / causal interpretation, not by spatial adaptation.

Our results support the independence hypothesis (I). The simplest account explains all major observations: a preceding distractor increases latency, a simultaneous distractor induces a bias, combining both distractors leads to a summed effect of the two. This separation maps naturally onto known anatomical and functional dissociations in the gaze-orienting system: the fixation system controlling latency vs. the multisensory spatial map determining the goal (Meredith & Stein, 1996; Sparks & Mays, 1983; Van Wanrooij et al., 2009).

### Relation to causal inference

Although independent mechanisms may be supported by the findings, the results may also align with a causal inference perspective (Körding et al., 2007; Shams & Beierholm, 2022). In this framework, integration occurs only when auditory and visual events are judged to belong to the same source. Temporal coincidence is a dominant cue for the unity judgment: simultaneous distractors may be, due to a high probability of a common cause (C=1), leading to integration, producing a spatial bias. In contrast, long preceding distractors might infer low probability of a common cause (C=2), leading to segregation, and no bias. In this view, the independence we observe may reflect two separate consequences of the same causal-inference computation: (1) spatial bias only when C=1 is likely, and (2) reaction time increases due to lingering segregation effects when C=2 is inferred. Thus, independence and causal inference are not mutually exclusive.

## Conclusion

By systematically manipulating the timing and spatial congruency of two visual distractors, this study reveals a robust separation between the spatial and temporal influences of vision on sound localization. Simultaneous distractors bias the spatial perception of sounds; preceding distractors delay orienting responses. When combined, these influences sum linearly with minimal interaction. These findings highlight that audiovisual integration during naturalistic orienting involves multiple, partly independent mechanisms: a temporal system governing disengagement and movement initiation, and a spatial integration system that combines cues only when temporal structure supports a common cause.

## Data availability

The data have been deposited in the Donders Institute for Brain, Cognition and Behaviour Data Repository at https://doi.org/10.34973/3dyz-d470

## Acknowledgements

We thank Günter Windau, Ruurd Lof, and Stijn Martens for valuable technical assistance.

## Author contributions

F.R., J.O., and M.W. conceived of the experiment(s). F.R. collected data. F.R., N.H, M.W. analysed the data. F.R. wrote the initial draft of the manuscript. All authors edited the manuscript.

